# Inferring the history of gene copy number evolution

**DOI:** 10.1101/2025.08.21.671444

**Authors:** Moritz Otto, Thomas Wiehe

## Abstract

Gene duplication plays a crucial role in the adaptive evolution and diversification of organisms by creating extra copies of genes that can evolve new functions while preserving the original. Duplicated genes can become fixed in populations or appear as copy number variants. However, inferring and dating these duplication events from present-day data is challenging, as gene copy count distributions could result from either a few ancient duplication events or many recent ones. Sequence based phylogenetic reconstruction, an often seen practice, does not include the history of individuals and hence may result in inconsistencies, which may lead to misinterpretations. Here, we introduce a novel model for inferring gene copy number evolution, which describes gene duplication and their evolution over time through a random walk on a coalescent duplication network. This approach is solely based on copy number counts and hence independent of the inconsistencies of sequence based inferences. Backward in time we implement structured coalescent simulations, where we re-interprete ‘structure’ as ‘genealogical distance’ based on copy number counts. We apply this model to the NB-ARC domain counts of NLR genes in *A. thaliana* to infer the number and times of duplication events that have led to the present day copy number distribution.

## Introduction

Multicopy gene families play a fundamental role in the evolution and adaptation of organisms by providing genetic redundancy, which allows functional diversification. Gene duplication events give rise to additional copies of genes, which can either be retained as redundant sequences, undergo subfunctionalization by partitioning ancestral functions, or acquire entirely new roles through neofunctionalization [1, 2, 3].

The adaptive potential of gene duplication has been demonstrated across a wide range of taxa, including mammals, fish, and various plant species, and across a wide range of gene functions, including immune and stress response or digestion [4, 5, 6, 7, 8, 9, 10].

Recent studies have explored the evolution of gene copy numbers in plant immune receptor families, particularly nucleotide-binding leucine-rich repeat (NLR) genes, which play a crucial role in pathogen defense. Relying on the long-read pan-NLRome data of *A. thaliana*, conserved NB-ARC domains were used as an indirect measure of NLR gene copy number [11, 12].

However, given the present-day copy number distribution, it remains unclear whether the observed gene copies originated from a few ancient duplication events or multiple recent ones (see Figure 1). Accurate estimation of the number and timing of events is required, for instance, to test if a recent ecological shift is synchronized with gene copy number alteration. Besides standard sequence based phylogenetic inferences several new tools were developed to identify and align gene copies [13, 14, 15]. Nevertheless, these sequence-based analyses of multicopy gene families require a well-defined theoretical null model to accurately interpret patterns of variation and divergence. Without such a framework, it becomes difficult to distinguish between signatures of neutral evolution, selection, or non-adaptive processes such as gene conversion. For instance, the presence of high sequence similarity among gene copies could be misinterpreted as evidence of recent duplication, when in fact it may result from homogenization via concerted evolution. Thus, theoretical modeling is essential to avoid biased or misleading conclusions in the study of complex gene families.

**Figure 1.**
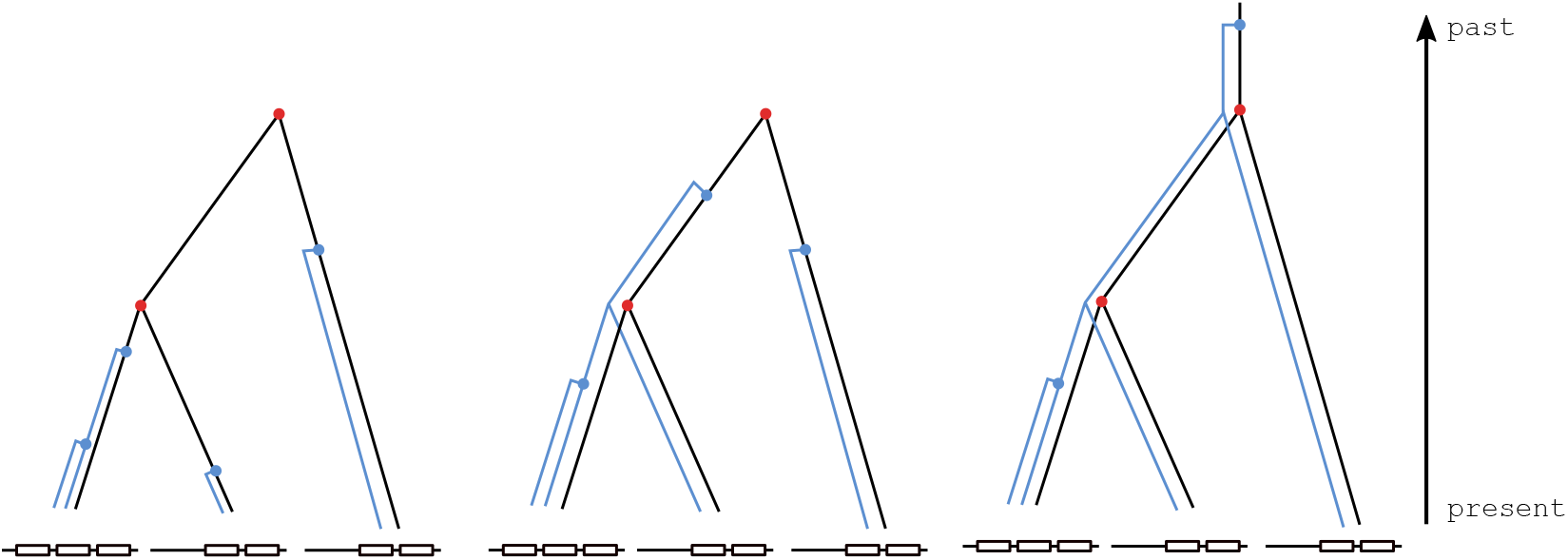
Example of three different duplication histories resulting in the same copy number distribution. Left shows four recent, independent duplications (blue dots and lines), center shows three duplications and right shows one ancient and one recent duplication. The different duplication events shape the tree topology, even though the coalescent events (red) are identical.

Here, we introduce a novel approach to model the history of multicopy gene families. Our approach starts from present day copy number counts and is twofold: First, we aim to estimate the most likely duplication rate that led to this present day configuration. This approach builds upon the work of [16], who utilized phase-type distributions in a random walk framework to estimate the joint site frequency spectrum in complex migration histories. We adapt this concept and introduce the **Coalescent-Duplication Network (CDN)**, see Figure 2. Starting with one single individual with one copy, the lineage may either split into two separate individual lineages (offspring event, red arrows), or into two linked copies within the same individual (duplication event, blue arrows). Following this process a too high duplication rate may lead to a copy number configuration that is not compatible with the present day sample anymore. Vice versa, a random walk with a too low duplication rate consists of too many individual lineage splits and is also not compatible with the present day distribution (gray arrows). Conditioning on reaching the desired copy number in the sample we calculate a maximum likelihood estimator of the duplication rate.

**Figure 2.**
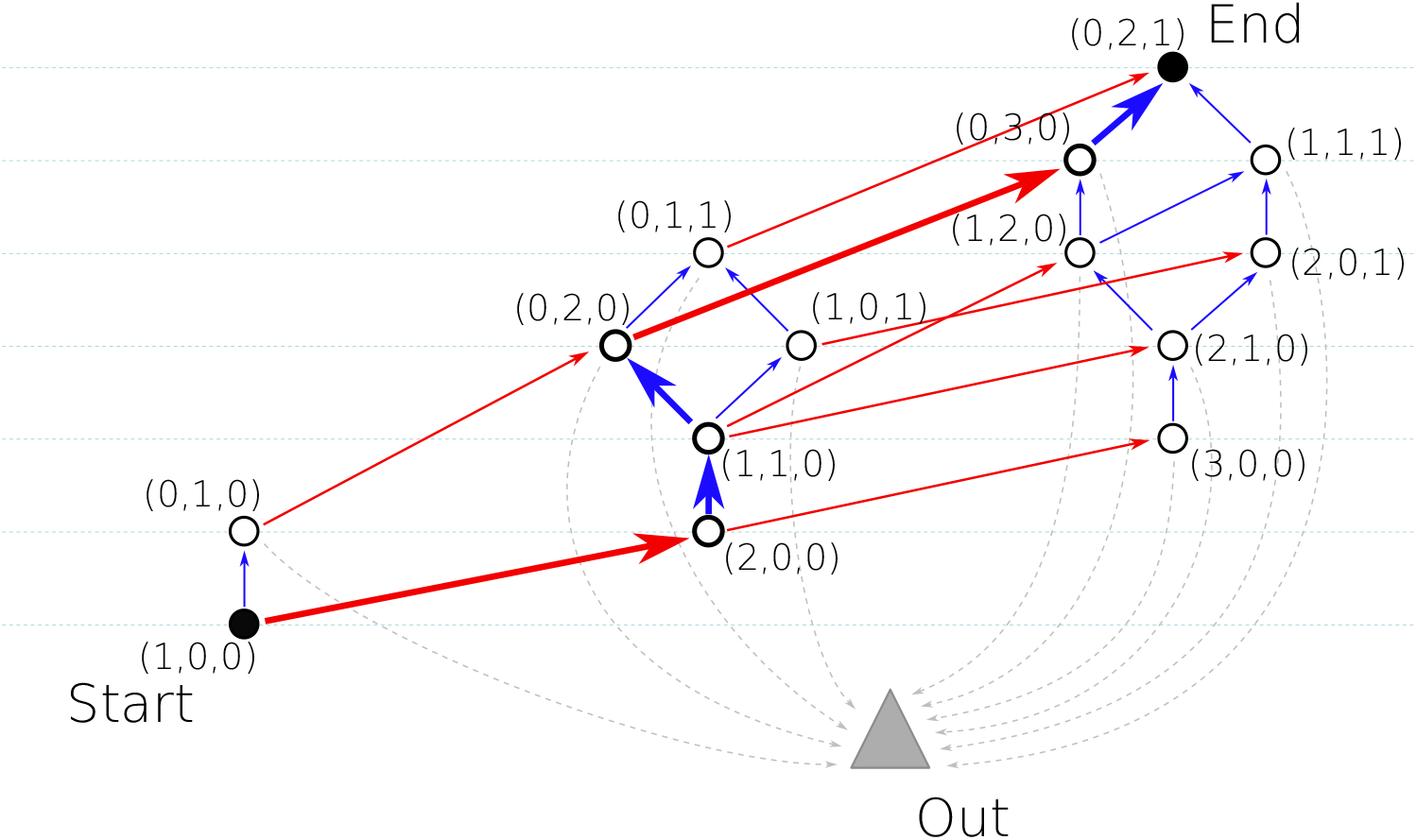
Example of a random walk on the Coalescence-Duplication-Network (CDN). Blue lines indicate duplication events, red lines split events. Duplication increases the copy number (dashed horizontal lines) by one, split events increase the number of individuals by one and may increase the number of copy numbers by one or more. Grey dashed lines indicate combined probabilities to reach a state, from which *α*_end_ is no longer possible and the random walk gets out-of-bounce. The path in bold lines corresponds to the central genealogy in Figure 1

In a second step, using this estimator, we simulate the backward-in-time process of this sample back to its initial single individual with a single copy in a coalescent framework. More precisely, we re-interprete the term of ‘structure’ (which is often used to describe geographic distance between individuals) as ‘genealogical distance’ in the sense of copy number differences, as already established in [17]. Therefore, two individuals may only be offspring of the same ancestor, if their copy number coincides. This intuitive assumption is currently ignored in sequenced based phylogenetic reconstruction and hence might be error-prone in inferring the evolutionary history (see Figure 3**A**, example of *A. thaliana*). Especially when considering effects such as selection or gene conversion acting on sequences, those inconsistencies of gene trees and individual trees might be misleading in downstream analyses. In contrast, our approach starts from simply counting the copy numbers of a given gene family in different individuals and traces back the most likely path to their ancestor. There are two ways in which genealogical lineages can merge: by coalescent events (red circles in Figure 3**B**), where lineages of different individuals meet, and by ‘de-duplications’ (blue circles), where copy count is reduced by one when looking backward in time.

**Figure 3.**
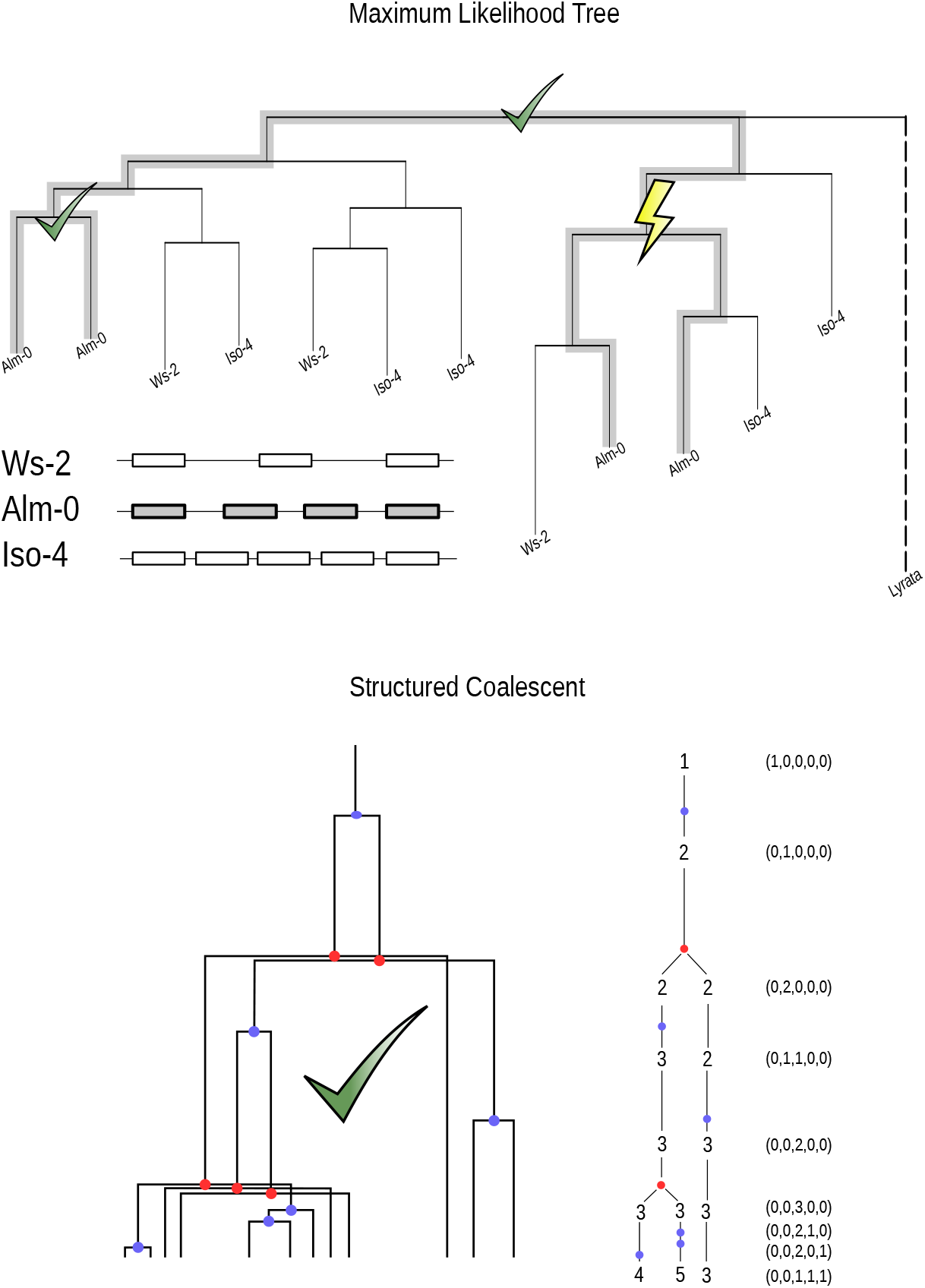
**A** Maximum likelyhood neighbour joining tree of **B3** NB-ARC domains in three accessions of *A. thaliana*, with the single copy of *A. lyrata* as outgroup (dashed line). Green shaded lines correspond to the lineages of the four copies of **Alm-0** accession. Intuitively, the common ancestor of two copies of the same individual has to be a duplication event. However, since duplication events affect the same lineage, the subtrees of a duplication event have to include the same individuals. Green marks mark the events, where this is true, whereas the lightning highlights the inconsistency of this phylogenetic tree. **B** Example of a structured coalescent simulation of the same copy number sample (3,4,5). By definition, this tree does not contain any inconsistencies. Blue dots indicate duplication events, red dots coalescent events. Note, that a coalescent event may merge more than two lineages if the individuals have more than one copy.

We implemented this structured coalescent framework to be compatible with the **msprime** and **tskit** packages [18]. Therefore, we are also able to superimpose the mutation process on the generated trees and to examine how duplication rates influence tree topology and, consequently, the estimation of key population genetic statistics such as Tajima’s *D* [19].

Finally, we applied our approach to a subset of 15 NLR-gene families of 10 *A*. thaliana accessions, with two families analyzed in more detail. We followed the workflow established by [11] to identify and align these gene copies. Using the CDN, we estimated the most likely duplication rate and, following the structured coalescent, we timed the duplication events and computed expected Tajima’s *D* under our model [19]. Our findings indicate that the most likely duplication history does not necessarily coincide with the most parsimonious one and that old duplications may lead to a biased Tajima’s *D*.

## Materials and Methods

### NB-ARC domains in A. thaliana

We followed the workflow of [11] to analyze the pan-NLRome^2^ [12]. Using the NB-ARC domains as proxies, we blast the annotated reference genes from the reference accession Col-0 (TAIR10.1 assembly^3^) against the pan-NLRome dataset. We were able to reproduce the copy number counts from [11] and align the corresponding gene families. For **B3**, which is assumed to be a young gene family, as it has only one copy in *A. lyrata*, we also built neighbour joining trees (Figure 3**A** and Figure 4). To test our model, we subsampled 10 accessions that showed a trustworthy alignment and represents the geographic distribution and the overall copy number distribution of two gene families (**B3** and **RPS5**), see Table S1 and Figure S1. Furthermore, we used the copy number counts of further NLR-genes from [11], scaled down to 10 individuals, as a test set to evaluate the duplication rate estimation procedure, see Table 1 and Figure 5.

**Table 1:**
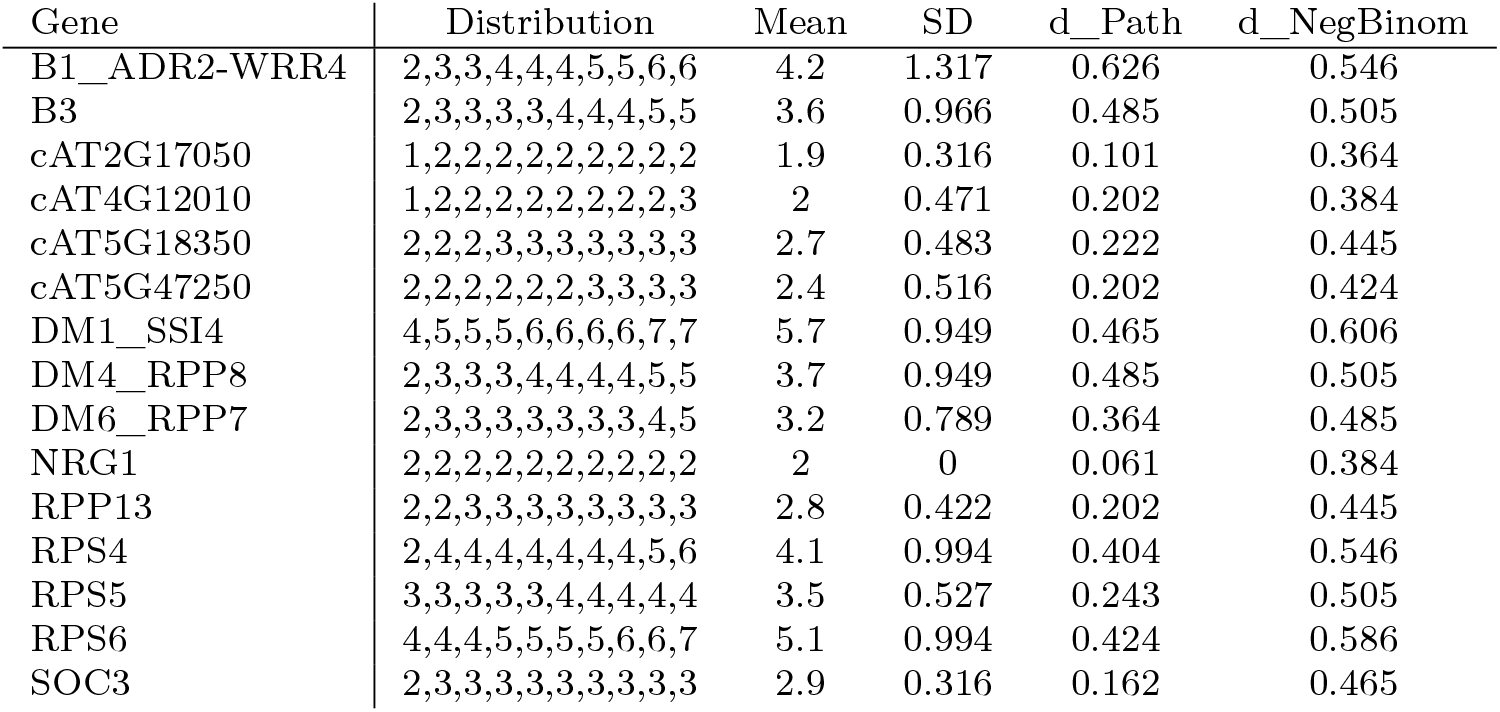
Duplication rate estimation for different distribution examples.

**Figure 4.**
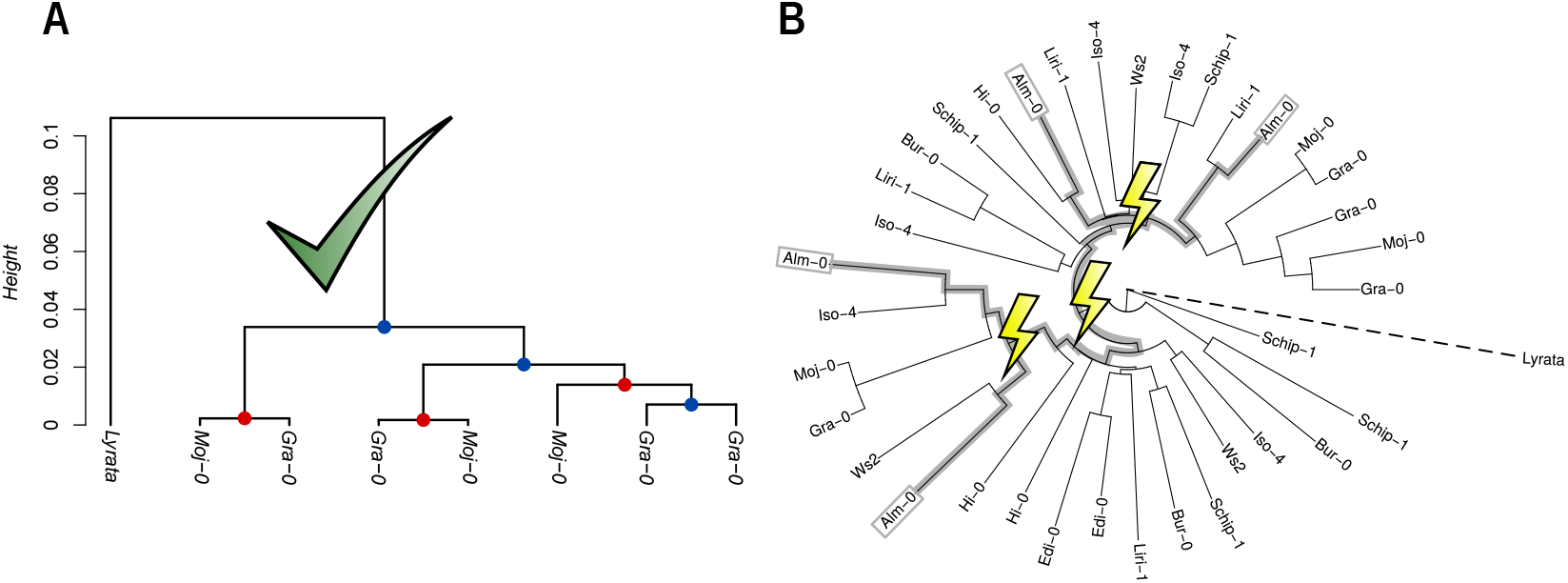
**A** UPGMA tree of **B3** sequences obtained from **Moj-0, Gra-0** and *A. lyrata* as outgroup. This shows a tree, which is consistent with the expected history of the two individuals, and hence we can place duplication events (blue dots) and coalescent events (red dots). **B** Radial neighbour joining tree for the sequences of **B3**, taken from 10 accessions, see Table S1 and Figure S1. Each leaf corresponds to a copy of **B3**. Grey bold lines show the genealogy of the four copies of **Alm-0**. Their ancestor node is assumed to be a duplication event. Lightnings mark inconsistencies of these duplication events, if the subtree leafs do not coincide (here in all three duplication events).

**Figure 5.**
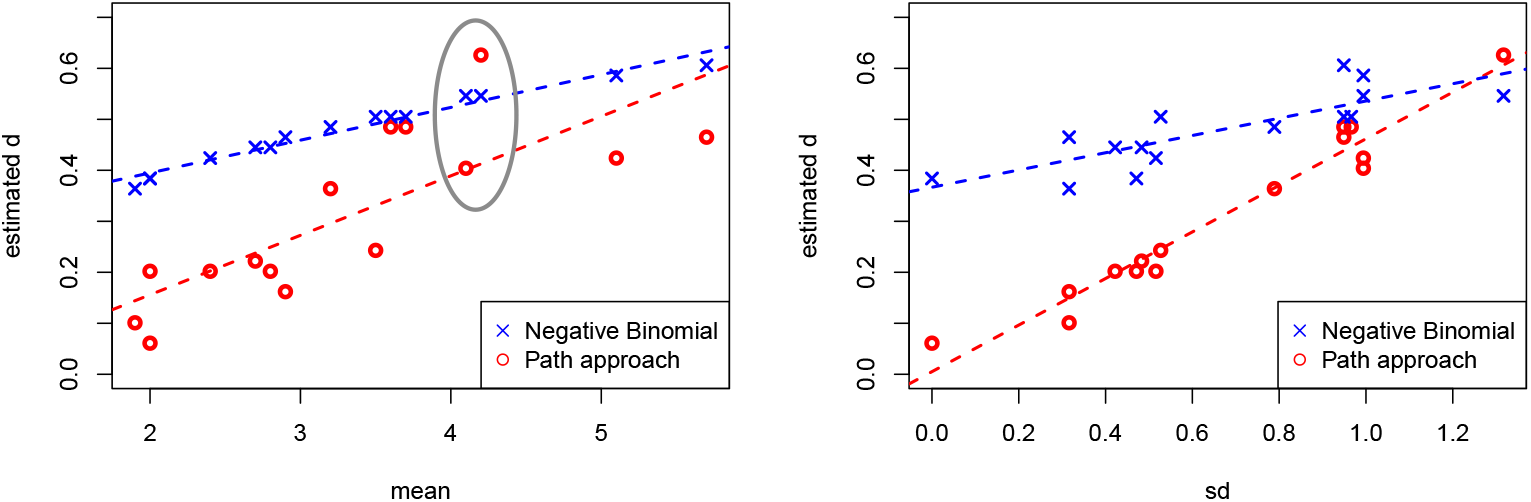
Correlation of duplication estimators *d*_path_, *d*_negbinom_ (scaled with *N*) with mean and standard deviation of copy number distributions from NLR genes, see Table 1. Grey circle shows estimates for **B3** and **RPS5**.

Note, that we do not aim to reproduce the data analysis done by [11] or [12] and do not proceed with an extensive sequence analysis but focus on the pure copy number counts as input data for our model.

### Duplication rate estimation

We encode the copy number distribution of a gene by a vector *α* = (*α*_*i*_)_*i*_, where *α*_*i*_ determines the number of individuals that have *i* copies. As an example, a sample of three individuals with (2,2,3) copies (as in Figure 1) is encoded as *α* = (0, 2, 1). Hence, thinking forward in time, we start with one single gene lineage in one individual, *α* = (1, 0, …). In each time step, a gene may duplicate with probability *d* and a lineage may split with probability 1*/N*, where *N* denotes the population size. Let **1**[*i*] be a vector with zeroes everywhere, except for a single 1 at position *i*. Then

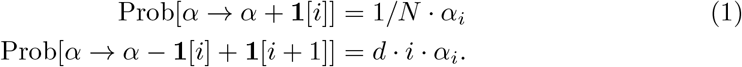

This results in a transient random walk in an infinitely large state space. However, we aim to calculate the probability that this random walk hits a certain state, i.e. our present day configuration. Since we only allow an increase of copy number or number of individuals, the random walk may reach a state from which it is impossible to end up in the desired state, *α*_end_. We summarize all these nodes as a cemetery state, which we simply call *α*_out_, since one steps out of the network without return. An example of *α*_end_ = (0, 2, 1) is shown in Figure 2.

Hence, the random walk either ends in *α*_out_ or *α*_end_. Using the Markov property, we denote the probability to start from *α* and reach *α*_end_ before hitting *p*_out_ as *p*_*α*_, where 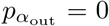 and 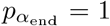. Therefore, we can determine the probability to start in *α*_start_ = (1, 0, …) and reach *α*_end_ by solving the linear equation system, where for all *α* we find

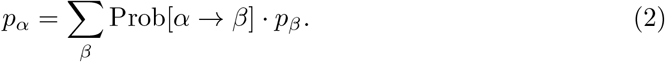

Therefore, given any *α*_end_ we can construct the CDN and apply the transition probabilities according to *d* and *N* and solve the linear equation system for 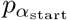 to determine the probability to generate the given copy number distribution, without hitting *α*_out_. This probability depends on *d* and *N*, and therefore we can use a maximum likelihood estimator to determine the duplication rate *d* that generates the copy number distribution.

Thinking differently, one may model duplication events similar to mutation events on a given coalescent tree, as done in [20]. Hence, one can think of the duplication process as a pure Yule-birth process. Therefore, the number of duplication events that have happened since the MRCA follows a negative binomial distribution which simplifies to a geometric distribution when assuming the ancestral genotype to be a single copy gene. More precisely, given a fixed time *T*, the first duplication event occurs at rate *d*, the second at rate 2*d* and so on. We define *X*_1_, *X*_2_, … as independent random variables with *X*_*i*_ ~ Exp(*i · d*) as the times between event *i −* 1 and *i* and 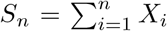 as the cumulative time of *n* events. Then, the number of events in the interval *T*, which we define as *Z*_*T*_, follows

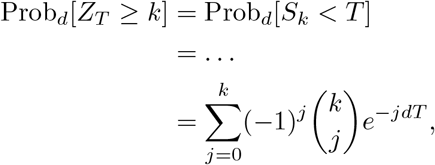

and hence *Z*_*T*_ ~ Geo(*e*^*−dT*^). Therefore, with *T* being the time until the most recent common ancestor, we find

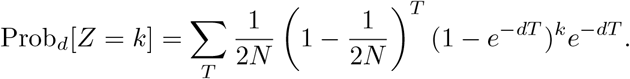

Concluding, we find two different approaches to get a maximum likelihood estimator of the duplication rate, *d*_path_ and *d*_negbinom_.

### Structured Coalescent

When simulating backward in time coalescence of multicopy gene families it is crucial to ensure that two individuals only coalesce, if both have the same number of gene copies. If two individuals coalesce, all their copies are paired, which leads to a merging of multiple lineages. Therefore, the process can be seen as a simulation of a backward in time random walk in the graph of Figure 2, using the previously estimated optimal duplication rate *d*. More precisely, starting with a copy number configuration given by *α*_end_ we include the waiting time spent in each state until we reach the most recent common ancestor of all gene copies, *α*_start_ = (1, 0, 0, …). We use the structure of **tskit** used in **msprime** [18] to implement a structured coalescent simulation in python. We trace the copy number distribution of a given sample and generate a random walk in the graph backward in time. Starting with *α* = (0, 1, 4, 3, 2) for **B3** and (0, 0, 5, 5) for **RPS5**, we ran 5.000 structured coalescent simulations for both copy number distributions. We count the number of duplication events in each random walk as well as the times of the duplication events measured relative to the TMRCA of individuals, to date the most likely duplication events. Furthermore, we uniformly place mutations on the tree branches according to their length with constant rate *µ*, as shown in Figure 3, to determine the effect of the duplication process on the tree topology.

## Results and Discussion

With the workflow established in [11] we were able to confirm their copy number counts for selected NLR genes. Two gene families, **B3** and **RPS5**, showed most reliable results. Furthermore, they both show same mean copy number of 4, but differ in their variance. Hence, their copy number counts were used as examples for further simulations. First, we focus on the results of the standard sequence based methods of phylogenetic inference. In Figure 4**A** we find a well-fitting UPGMA gene tree for **B3** in the accessions of **Moj-0** and **Gra-0. Gra-0** experienced one recent duplication event (blue dot), before coalescing with **Moj-0**, and indeed, all the coalescent events of the three copies occurred at the same time (red dots). However, this consistency vanishes, when building the alignment and the neighbour-joining tree of all ten accessions. Considering the same example of **Alm-0** as in Figure 3, we find inconsistencies in this NJ tree. More precisely, if one traces back the history of two copies of the same individual, their common ancestor has to be a duplication event. Moreover, duplication events affect the same lineage and therefore the subtrees of the duplication event have to include the same individuals. Intuitively speaking, if all individuals of a sample have two copies and this duplication event was ancient, one would find two subtrees with all the individuals as leafs. However, in the example of the four copies of **B3** in **Alm-0**, none of the three duplication events satisfies this condition (marked as lightnings in Figure 4**B**). More strikingly, each of the four gene copies of **Alm-0** is paired with a copy from a different accession.

The chosen gene families **B3** and **RPS5** both have a mean copy number of ~ 4, but differ in their variance. For both distributions we calculated 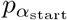 in the CDN for various *d*_path_*N* in (0,2). The widespread distribution of **B3** led to an estimation of *d*_path_*N ~* 0.469, around twice as high as the one of **RPS5** (Figure 6). This difference is biologically plausible, since a low-variance distribution may result from only a few older duplication events that have since stabilized, whereas a broader (high-variance) distribution suggests a greater number of more recent duplications contributing to ongoing diversification. Indeed, we obtain this result for the chosen copy number distributions of NLR genes of [11] (see Figure 5 and Table 1). When comparing *d*_path_ with *d*_negbinom_ we observe that both correlate with the mean and the standard deviation of the given copy number distribution. However, *d*_path_ is more sensitive towards the variance, whereas *d*_negbinom_ aligns better with the mean. Furthermore, *d*_path_ tends to estimate a lower duplication rate and has a larger range of estimates (0.1 - 0.6) compared with *d*_negbinom_ (0.4 - 0.6). This highlights the strength of the new estimator, as it includes the information of the variance of the distribution. Conceptually, a high mean copy number is not neccesarily an indicator of a high duplication rate, if the variance of the distribution is low. This distribution might rather be a result of a few, ancient duplications. The *d*_negbinom_ does not cover these differences, as indicated in the example of **B3** and **RPS5**, which have the same mean value and result in the same *d*_negbinom_, despite their different variance (grey circle in Figure 5).

**Figure 6.**
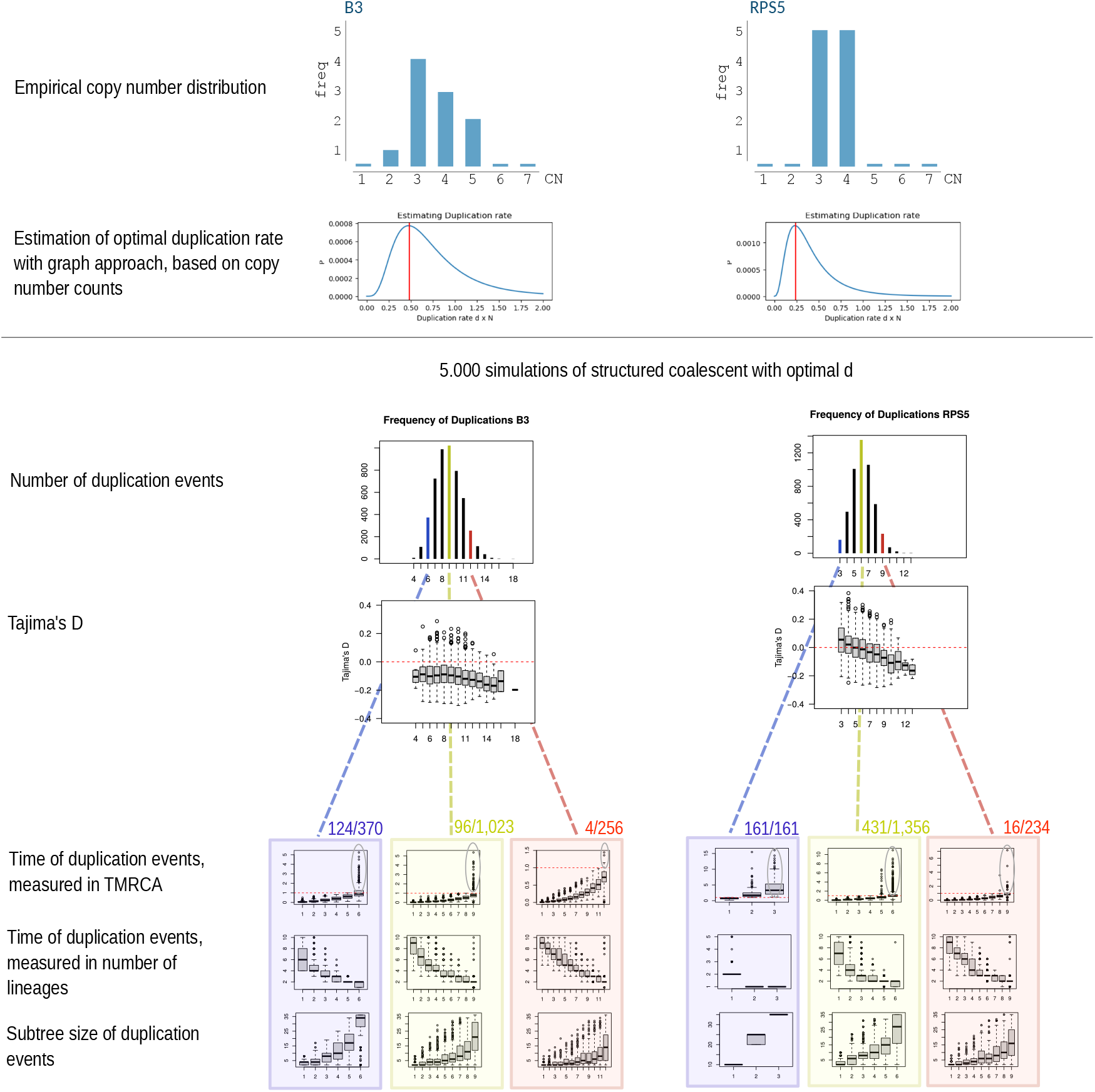
Estimation of the duplication rate for two examples (**B3** and **RPS5**) of gene copy number distribution. Top shows copy number distribution. Second row shows 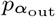 for duplication rates *dN* ranging from 0 to 2, with vertical red line indicating maximum likelihood. Third row shows frequency table of duplication events for 5,000 backward in time structured coalescent simulations. Yellow bar highlights the most frequent one. Fourth row shows Tajima’s D given the number of duplications. Bottom boxplots show the time and the subtree size of the duplication events. Ancient duplication events (i.e. that occurred before the TMRCA) are highlighted with gray circle and the corresponding numbers.

Given these duplication rate estimators, we aim to infer the number of duplication events leading to this distribution and to date those events. Therefore, we conducted 5,000 structured coalescent simulations for parameter sets *α* = (0, 1, 4, 3, 2) with *dN* = 0.469 (**B3**) and *α* = (0, 0, 5, 5) with *dN* = 0.224 (**RPS5**). For **B3**, a total of 1,023 simulations (20%) resulted in exactly nine duplications, representing the most frequent and thus most likely scenario (see Figure 6). In the case of **RPS5**, 1,355 simulations (27%) featured six such events. Recap that these results were obtained solely by the copy number counts, independent from sequence data. The results suggest that the observed copy number distributions are unlikely to arise from either a few ancient duplications or an accumulation of very recent ones alone. When focusing on the temporal distribution of the most likely number of events (Figure 6, yellow boxes), the inferred timings continuously span up to the most recent common ancestor (indicated by the red dashed line), which is consistent with a model that assumes constant and ongoing duplication pressure. For the subset of 1,023 **B3** simulations yielding nine duplications, only 96 trajectories included an event predating the most recent common ancestor of the sample. The remaining runs indicate that the duplications happened relatively recently. This strongly supports a scenario in which the observed copy number variation arose from approximately nine independent and recent duplication events. In contrast, for **RPS5**, a minimum of three duplications is required to reproduce the empirical copy number distribution. With *dN* = 0.24, only 161 of the 5,000 simulations met this minimum. The most frequently observed scenario comprises six duplications, where 431 out of 1,355 simulations involved at least one duplication predating the most recent common ancestor. This outcome is notable, given that the observed bimodal distribution – three and four copies among equal proportions of the sample – would intuitively suggest two ancestral duplications followed by a more recent one affecting only part of the population. However, under the assumption of a constant and uniformly acting duplication rate, our model instead favors a scenario involving six independent events distributed over time.

Note, that the structured coalescent simulations strongly depend on the correct estimation of *d*, since the transition probabilities of the backward in time random walk on the graph of Figure 2 depend on the duplication rate. Hence, with a too high duplication rate one expects several duplication events to occur recently, i.e. starting at *α*_end_ the random walk always follows the next blue path in Figure 2. But then, no further ‘deduplication’ events are possible, which means that – even though we assume a constant duplication force – there will be no duplication event on the long branches in the coalescent tree. Vice versa, a too low duplication rate assumes that all duplication events have happened before the most recent common ancestor of the individuals, leading to long branches before the MRCA. Therefore, both a too high and a too low *d* estimate shape the topology of the coalescent tree and hence affect population statistics.

With maximum likelihood estimated *dN*, we placed 500 mutations on the outcoming tree for each of the 5,000 simulations and calculated Tajima’s *D* (see Figure 6). We observe a small bias towards a negative *D* in **B3**, which appears to be negligible. Therefore, with correct estimator and constant duplication rate, the tree topology appears to be similar to that of the neutral coalescent. Note, that this simulation scheme takes into account both coalescent of individuals and de-duplications of copies and results in a genealogical tree of gene copies.

## Caveats and Outlook

Here we presented a novel approach to estimate the duplication rate of multicopy gene families based on their copy number counts. We implement the evolution as a random walk on the **Coalescent Duplication Network**, where the transitions from one state to another correspond to either duplication or coalescent events. However, we had to limit our estimation procedure on intermediate copy number counts and samples of size *n* = 10, as the graph network increases exponentially in size with additional copies and additional sampled individuals. Solving this scaling problem and subsequent analysis of the CDN needs to be addressed in the future. With larger sample sizes and larger copy numbers we also aim to include larger sequence analyses, including information of the population structure. Implementing selective pressure on gene copy number and, in the case of immune genes, diversifying selection acting on gene function as done in [21, 22] may represent another prospective goal.

Concluding, our results show the necessity of a duplication model when analyzing multicopy gene families and highlight the limitations of standard tree inference approaches and summary statistics when there is no clear distinction between paralogs and orthologs.

## Data Availability Statement

All data and custom codes used for simulations and analysis can be found at https://github.com/Moritz-Otto/motto-randomwalk

## Supplementary Material

**Table S1:** Subsample of 10 accessions that showed a trustworthy alignment and represents the geographic distribution and the overall copy number distribution of the two gene families.

| AccID | AccName | relict | MAGIC | latitude | longitude | B3-CN | RPS5-CN |
| --- | --- | --- | --- | --- | --- | --- | --- |
| 6981 | Ws-2 | 0 | 0 | 52.3 | 30 | 3 | 4 |
| 7058 | Bur-0 | 0 | 1 | 54.1 | -6.2 | 3 | 3 |
| 7111 | Edi-0 | 0 | 1 | 55.9494 | -3.16028 | 2 | 3 |
| 7167 | Hi-0 | 0 | 1 | 52 | 5 | 3 | 4 |
| 9518 | Alm-0 | 1 | 0 | 39.8848 | -0.364008 | 4 | 4 |
| 9543 | Gra-0 | 1 | 0 | 36.7691 | -5.39373 | 4 | 3 |
| 9550 | Iso-4 | 1 | 0 | 43.049 | -5.37103 | 5 | 3 |
| 9654 | Liri-1 | 0 | 0 | 41.413538 | 13.771045 | 4 | 4 |
| 9721 | Schip-1 | 0 | 0 | 42.72 | 25.33 | 5 | 3 |
| 9869 | Moj-0 | 1 | 0 | 36.76 | -5.28 | 3 | 4 |

**Figure S1:**
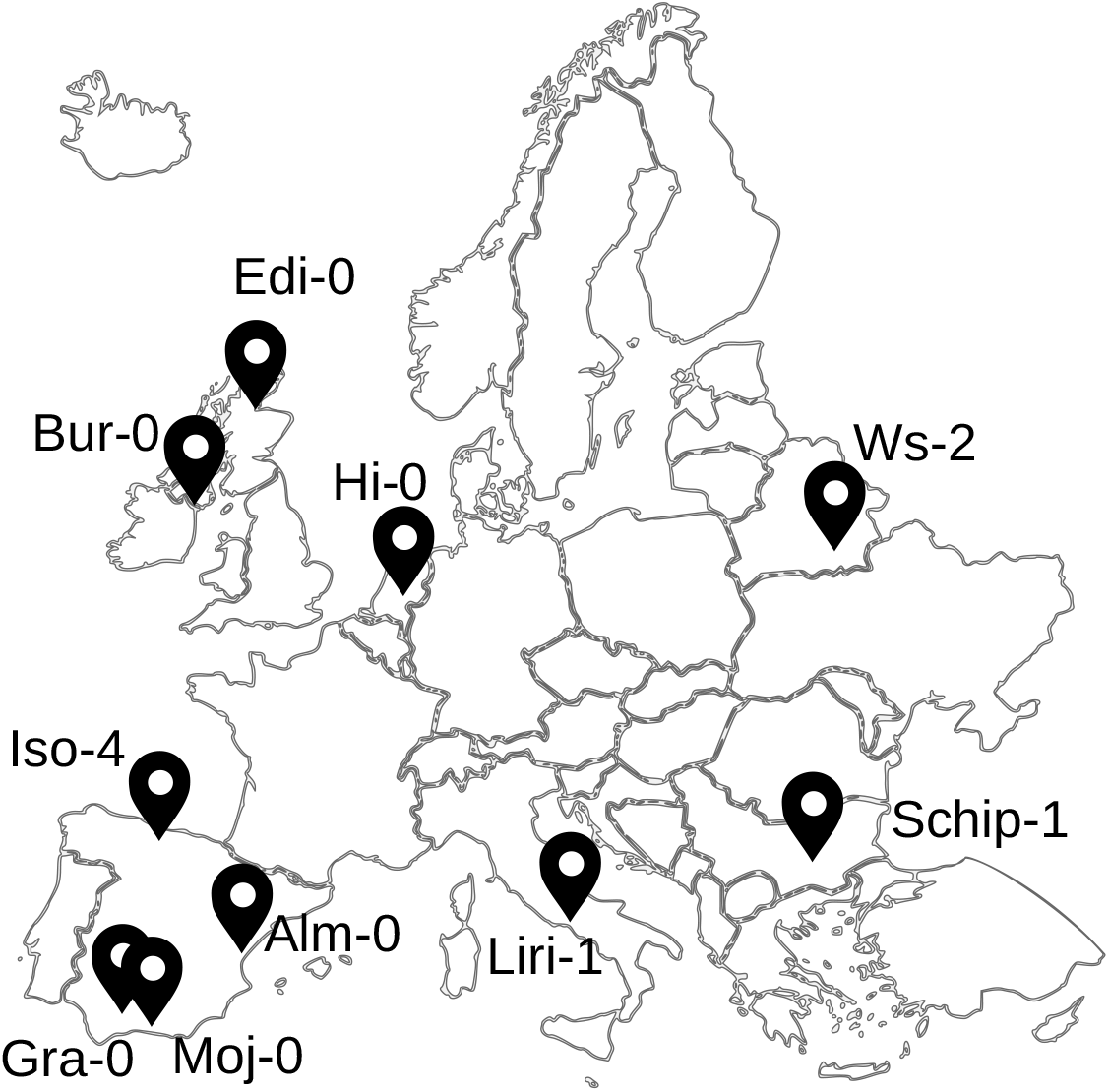
Subsample of accessions selected from [12]

## Footnotes

1 http://ftp.tuebingen.mpg.de/ebio/alkeller/pan_NLRome/

2 https://www.ncbi.nlm.nih.gov/datasets/genome/GCF_000001735.4/

