## Supplementary figures and images for "Inferring the history of gene copy number evolution"

### Supplementary Figure 1

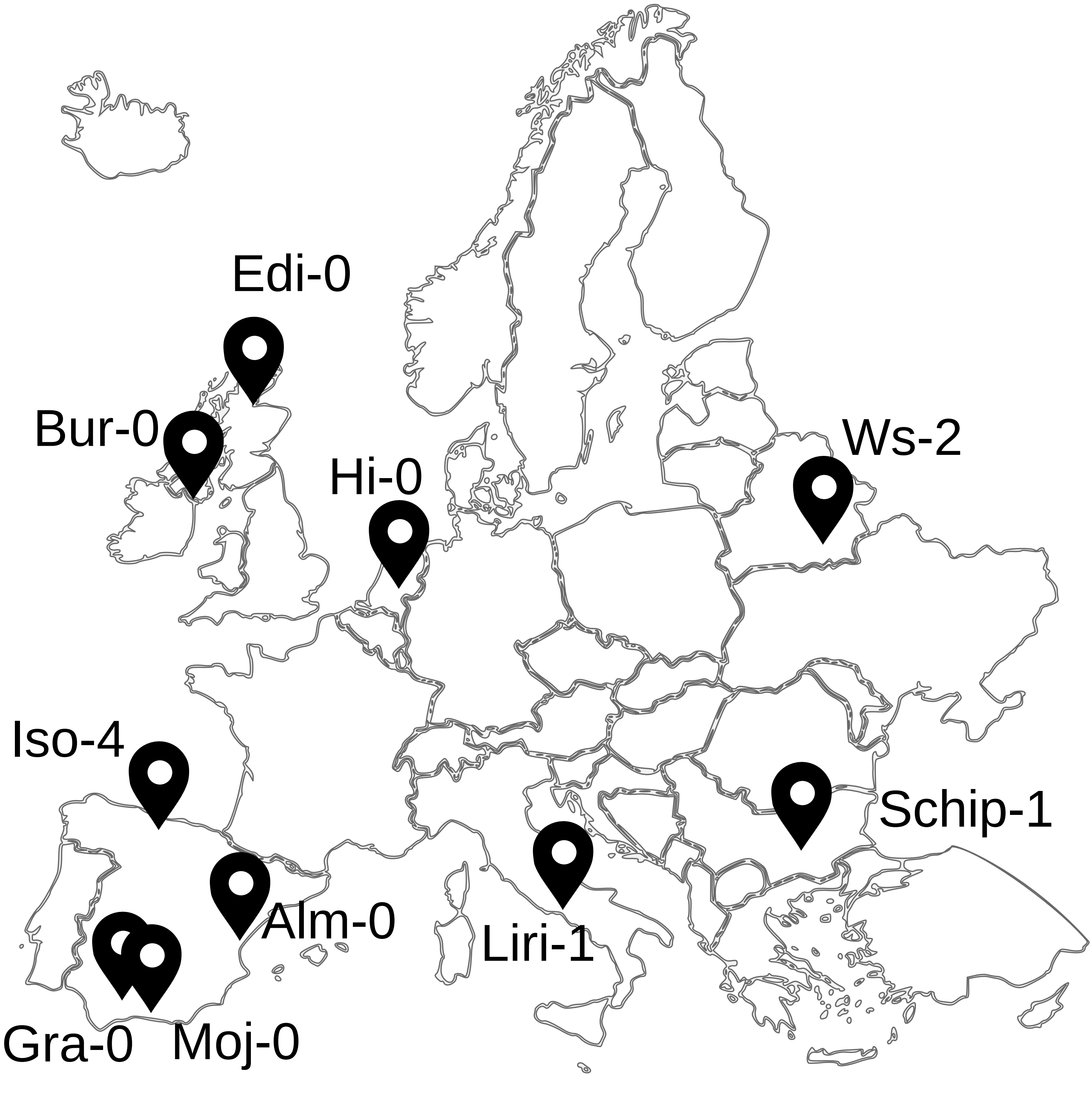
